# Transcriptomic Insights into the Virulence of Acinetobacter baumannii During Infection: Role of Iron Uptake and Siderophore Production Genes

**DOI:** 10.1101/2024.07.15.603485

**Authors:** Kah Ern Ten, Sadequr Rahman, Hock Siew Tan

## Abstract

*Acinetobacter baumannii* is a top-priority pathogen as classified by the World Health Organisation. It causes life-threatening infections in immunocompromised patients, resulting in prolonged hospitalisation and high mortality. Increasing cases of community-acquired *A. baumannii* infections with rapid progression and severe infections have been reported. This study used the previously described *Galleria mellonella* infection model to investigate the virulence mechanisms of the community strain C98 (Ab-C98) via transcriptomic analysis using direct RNA sequencing. This strain showed greater killing and more rapid colonisation in the larvae than a clinical reference strain (ATCC BAA1605). Differential gene expression analysis revealed the significant upregulation of three major iron clusters: the acinetobactin and baumannoferrin clusters for siderophore production and the Feo system for ferrous iron uptake. Targeted knockout of siderophore production genes (*basC*, *bfnD* and isochorismatase family protein) significantly attenuated virulence in mutants with minimal impact on the bacterial growth *in vivo*. Overall, this study highlights the virulence of *basC*, isochorismatase family protein and *bfnD* in the pathogenicity of *A. baumannii*. As these targets are highly conserved in *A. baumannii* and the closely related *A. pittii* and *A. lactucae*, they could serve as potential therapeutic targets for developing new antivirulence agents to combat these pathogens.

## 1. Introduction

*Acinetobacter baumannii* is the most clinically significant member of the *A. calcoaceticus*-*baumannii* (Acb) complex [1]. This bacterium is known as a nosocomial pathogen due to its frequent isolation from hospital settings. With the ability to develop drug resistance rapidly and persistence on dry surfaces, this pathogenic bacterium poses a great risk to hospitalised immunocompromised patients, causing prolonged hospitalisation and high mortality [2]. Previous research studies have used clinical strains for antimicrobial resistance and virulence studies to understand the pathogenesis of this bacterium [3–8]. Although community-acquired

*A. baumannii* contributed to a minor proportion of *A. baumannii* infections, increasing cases of community-acquired *A. baumannii* in tropical and subtropical regions have also been reported [9–14].

Iron is an essential micronutrient for most living organisms. It is required for many major biological processes, such as DNA biosynthesis, oxygen transport, the trichloroacetic acid (TCA) cycle and regulation of gene expression [15,16]. Despite its essentiality, iron can catalyse reactions generating free radicals that cause damage to the membrane components, nucleic acids and other essential biological materials [17]. Therefore, the iron content is regulated by stringent iron homeostasis to control the potentially toxic effects of iron. During a pathogenic bacterial attack, the host restricts the iron availability in circulation by storing the iron intracellularly due to immune responses against invading pathogens [18]. Thus, the iron acquisition mechanisms represent crucial virulence factors the pathogenic bacterium employs to thrive within a host. *A. baumannii* possesses an arsenal of intrinsic virulence factors. For example, to overcome the iron-restricted condition in the host environment, *A. baumannii* can invade the intracellular iron-containing host cells to release iron by expressing phospholipases [19] and uptake the iron via its multiple iron acquisition systems. Siderophores are high-affinity iron-chelating molecules that aid the bacterium in scavenging iron from the host’s iron-protein complex [20]. The most common siderophore systems identified in *A. baumannii* include acinetobactin, baumannoferrin and fimsbactin [21]. Other iron transport systems, such as the TonB-ExbB-ExbD energy-transducing complex for the active transport of ferric-siderophores or the Feo system for active uptake of ferrous iron, have also played significant roles in *A. baumannii* under iron-limiting conditions [22]. Targeting the iron acquisition mechanism showed the potential to reduce bacterial pathogenesis, as the disruption of this system markedly decreases the ability of the pathogens to colonise the host [23]. Recently, gene expression profiling of *A. baumannii* during infection in a host has been performed in murine models, and iron acquisition pathways were enriched in *A. baumannii* in the host environment [24–26]. However, these research studies were mostly performed on *A. baumannii* clinical isolates or ATCC reference strains, and there is limited knowledge of the virulence factors of community-acquired *A. baumannii* and how they behave in a non-mammalian model. We hypothesised that the rapid progression of diseases by community-acquired *A. baumannii* could be due to their intrinsic virulence factors similar to hospital strain. Due to the increased usage of non-mammalian models, it is also important to understand the *A. baumannii* biology in these models to elucidate the bacterial pathogenesis better.

In this study, we have used *Galleria mellonella*, an insect model in which we had previously established a protocol for *A. baumannii* virulence studies using the community-acquired *A. baumannii* strain C98 (Ab-C98) as the infection pathogen, which caused a high mortality rate in *G. mellonella* [27]. In this study, we compared the killing of the clinical reference strain ATCC BAA1605 in the *G. mellonella* model; however, the clinical strain showed lower killing than the Ab-C98. Therefore, to identify the virulence factors of the Ab-C98, we performed a transcriptomic analysis of this bacterium by employing the Direct RNA sequencing technique (Oxford Nanopore Technologies), which allowed us to identify potential virulence targets that could help in the development of new antimicrobials against this pathogen.

## 2. Methods

### 2.1 Strains and growth conditions

The wild-type *A. baumannii* strains used in this study, strain C98 and ATCC BAA1605, were kindly gifted by Nazmul Hasan Muzahid (Monash University Malaysia). Wild-type *A. baumannii* strains were routinely cultured on Leeds *Acinetobacter* agar (Himedia). Isogenic mutant derivatives were maintained on Luria-Bertani agar (Himedia) supplemented with 50 μg/mL apramycin. All bacterial strains were grown aerobically in LB broth (Himedia) at 37 °C, with shaking at 200 rpm, to stationary phase conditions (∼20 h) unless stated otherwise. The insect model, *G. mellonella*, was maintained at room temperature at the 6th instar stage and fed with artificial food as we previously established in Ten, Muzahid [27]. Any melanised or inactive larvae were discarded from the experiment.

### 2.2 *G. mellonella* infection assays for wild-type and isogenic mutant derivatives

The virulence of *A. baumannii* strains was tested by performing a *G. mellonella* killing assay and bacterial burden assay as described previously [27]. The bacterial suspension with appropriate cell density was injected into the last left proleg of the healthy, non-melanised 6^th^ instar stage larvae and incubated at 37 °C in a standard bacterial incubator. Larvae were considered dead when they were unresponsive to physical stimuli and melanised. The experiments ceased when pupation occurred. Larvae injected with only sterile PBS and larvae without receiving any injections were used as the control groups. Ten larvae were used for each experimental group, and all experiments were carried out in three independent replicates (n=30). Only the experiments where all non-manipulated larvae survived throughout the experiment were included in the analysis. The extracted hemolymph with appropriate dilution was plated onto Leeds Acinetobacter agar plates for the wild-type strain or agar plates supplemented with 50 μg/mL apramycin for the mutant derivatives to quantify the bacterial load. Three independent replicates were carried out. Larvae injected with PBS only were used as the negative control. Only the experiments with no colonies obtained from the PBS-injected control were used for analysis.

### 2.3 Biological RNA sample collection

The bacterial RNA was isolated from the *in vitro* broth culture (no infection control, IVB) and the *in vivo* infected samples from infected *G. mellonella* larvae (infection, IVV) as described previously [27]. Briefly, the bacterial cells were harvested from the infected larvae at 3-h post-infection (p.i.), enriched, and the bacterial RNA was isolated. The mid-exponential phase *A. baumannii* grown in an LB medium was used for the no-infection control. The absence of gDNA in the extracted RNA samples was verified by PCR to amplify the housekeeping gene *rpoD*. The RNA integrity was assessed, and the absorbance values (A260/230 and A260/280) and the concentration of the isolated RNA were quantified via a Nanodrop spectrophotometer. The extracted RNA was stored at -80 ℃. Two biological replicates were collected for each infection and no-infection group.

### 2.4 RNA treatment and MinION Direct RNA sequencing

Before the nanopore sequencing, the RNA samples were poly-adenylated using *E. coli* poly(A) polymerase (NEB) [28]. Briefly, 5 μg of total RNA was incubated at 70 ℃ for 3 minutes to remove the secondary structure. Then, the RNA was mixed with 20 units poly(A) polymerase, 2 μL reaction buffer and 1 mM ATP and incubated for 15 min at 37°C in a total reaction volume of 50 μL. The RNA quality was assessed, and poly(A)-tails were confirmed by reverse transcription using oligo d(T)_23_VN and PCR amplification of housekeeping gene *rpoD*. The barcode sequences were synthesised by Integrated DNA Technologies (IDT) (Supplementary Table S1), and the sequencing libraries were barcoded following the conditions established by Smith and Ersavas [29]. The sequencing libraries were prepared using the Direct RNA sequencing kit (SQK-RNA002), following the protocol established by Oxford Nanopore Technologies (DRS_9080_v2_revO_14Aug2019). Briefly, 1.2 μg of poly(A)-tailed RNAs were ligated to pre-annealed custom RT adaptors (IDT) in the presence of RNA calibration strand (RCS) using concentrated T4 DNA Ligase (NEB). The ligated products were reverse transcribed using ProtoScript® II Reverse Transcriptase (NEB). The resulting RNA:cDNA hybrids were purified using 1.8× Agencourt RNAClean XP beads and washed with freshly prepared 70% ethanol (molecular grade). Then, 50 ng of reverse-transcribed RNA from each library was pooled and ligated with the RMX adapter comprising motor protein. The mix was then purified using 1× Agencourt RNAClean XP beads and washed with wash buffer twice. The sample was eluted in 20 μL of elution buffer and mixed with RNA running buffer, loaded onto a primed R9.4.1 flow cell (FLO-MIN106) and run on a MinION sequencer with MinKNOW acquisition software version 22.12.7. Samples were reloaded onto the nanopore flow cells, and the sequencing run was continued until no active pores were found.

### 2.5 Data analysis

#### 2.5.1 Nanopore sequencing data pre-processing

The fast5 files from the ‘pass’ and ‘fail’ folders from the two sequencing runs were basecalled using Guppy version 6.3.8 (Oxford Nanopore Technologies) using RNA-specific parameters (--calib_detect --reverse_sequence --u_substitution). Only the reads that passed the quality threshold (minimum qscore = 7) were written into the ‘pass’ folder and were used for subsequent analysis. DeePlexiCon demultiplexed the raw fast5 files [29], and the reads of each library were split according to their barcode numbers with a confidence score of above 0.95. The fastq files of each library were trimmed using Chopper version 0.2.0 [30] using parameters (-q 7 -l 200 --tailcrop 100). All libraries were aligned to the Ab-C98 whole-genome sequence retrieved from the NCBI database(https://www.ncbi.nlm.nih.gov/assembly/GCF_024357825.1) using Minimap2 version 2.24-r1122 [31] with noisy Nanopore Direct RNA-seq parameters (-ax splice -uf -k14 --secondary=no) [32]. For *in vivo* libraries, the nanopore reads were first mapped to the *G. mellonella* reference genome (https://www.ncbi.nlm.nih.gov/datasets/genome/GCF_026898425.1/). Then, the unmapped reads were extracted using SAMtools [33] and aligned to the Ab-C98 whole-genome sequence using the same parameters described above. Reads mapped to the Ab-C98 were converted to bam files, sorted and indexed using SAMtools.

#### 2.5.2 Differential Gene Expression Analysis

The changes in the gene expression levels of Ab-C98 during the infection were analysed by differential gene expression (DGE) analysis. The GenBank gene annotation files of Ab-C98 were retrieved from the NCBI database. The number of reads assigned with feature ‘CDS’ was quantified by featureCounts in long read mode (Galaxy Version 2.0.3+galaxy2). Then, DGE analysis was performed using DESeq2 (Galaxy Version 2.11.40.8+galaxy0). Only genes with adjusted *p*-value < 0.05 and log_2_fold-change of 1 were considered differentially expressed. Spearman correlation between the two biological replicates in each treatment group was assessed. A volcano plot was used to visualise the results of the DGE analysis and plotted by SRplot [34]. The heatmap of the VST normalised read counts of the differentially expressed genes (DEGs) was constructed using GraphPad Prism version 10.0.3. Functional enrichment analysis and functional association of genes were performed via the STRING version 12.0 [35] and plotted by SRplot. The gene locus of the acinetobactin, baumannoferrin and *feo* genes were plotted using gggenes (https://github.com/wilkox/gggenes.git). The promoter regions were predicted by the web tool BPROM (http://www.softberry.com/berry.phtml).

### 2.6 Reverse-transcription quantitative PCR

The expression levels of the shortlisted targets in the infection samples and the no-infection controls were assessed by RT-qPCR. The reverse transcription quantitative PCR (RT-qPCR) assays were performed using the CFX96 Real-Time System (BioRad). The primers used in this study are listed in Supplementary Table S2. Amplifications were carried out in 20 μl reaction solutions containing 1X Luna® Universal qPCR Master Mix (NEB), 20 ng cDNA and 0.25 μM each gene-specific primer, with the conditions according to the manufacturer’s instructions. The housekeeping gene *rpoD* was used as the reference gene. Each assay was performed with three technical replicates for each of the three biological replicates. The relative fold-change of the gene expression level compared to the reference gene (*rpoD*) between the two condition groups was calculated using the Pfaffl method [36].

### 2.7 Synthesis of donor template for gene knockout

Overlap extension PCR (OE-PCR) was performed to synthesise large chimeric DNA fragments carrying the selection marker (apramycin resistance gene, *apmR*) flanked by 300 bp-long fragments homologous to sequences flanking the target site. The oligonucleotides used for the donor template construction were summarised in Supplementary Table S2. Briefly, the homology arms of the target genes were amplified from the genomic DNA (gDNA) of Ab-C98 using Taq DNA Polymerase with Standard Taq Buffer (NEB), and the *apmR* was amplified from the plasmid pCasAb-apr [37]. All the PCR products were purified using Monarch PCR & DNA Cleanup Kit (NEB). Equimolar concentrations of the purified PCR products were mixed and subjected to OE-PCR using the following conditions: 95 °C for 30s, then 5 cycles of 95°C for 30s, 54.2 °C for 1 min and 68 °C for 1 min 25s without the addition of primers. Then the reaction was paused, and a final concentration of 0.2 μM of primers binding at the ends of the homology arms was added, followed by 35 cycles of 95 °C for 30s, 54.2 °C for 1 min and 68 °C for 1 min, 25s, lastly 68 °C for 5 mins and hold at 4 °C. The PCR products were purified and concentrated via CentriVap Benchtop Vacuum Concentrator to approximately 10 μL, yielding a final concentration of 1000 ng/μL.

### 2.8 Construction of isogenic mutant derivative strains by gene replacement

Gene knockout in Ab-C98 was conducted via the RecAb recombinase system, encoded by the plasmid pAT02 [38]. Ab-C98 harbouring plasmid pAT02 was grown in LB broth supplemented with ampicillin 100 μg/mL at 37 °C, 200 rpm for 18 hours. The culture was then diluted to OD_600nm_ = 0.05 in 50 mL LB broth and grown at 37 °C, 200 rpm. IPTG was added to a final concentration of 2 mM after 45 minutes of incubation, and the bacteria were harvested at OD_600nm_ = 0.7. After 3 washes in 25 mL ice-cold 10% glycerol, 100 μL of electrocompetent cells (∼10^10^ bacteria) were mixed with 5 µg – 10 µg of donor template and electroporated in a 2-mm cuvette at 2.5 kV. After outgrowth in 4 mL SOC medium containing 2 mM IPTG, the bacteria were pelleted, plated on 50 µg/mL apramycin LB agar, and incubated overnight at 37°C until colonies appeared. Successful knockout mutants were confirmed via colony PCR and routinely maintained in an LB medium supplemented with 50 µg/mL apramycin. Plasmid curing was performed by continuous passage of the Ab-C98 mutants in 1:1000 LB broth with apramycin, grown overnight, and streaked onto 50 μg/mL apramycin and 100 μg/mL ampicillin LB agar until no visible colonies on the ampicillin agar plate.

### 2.9 Growth rate analysis of knockout mutants under normal and iron-limiting conditions

Growth rates of the wild-type Ab-C98 and the mutant derivative strains were assessed by measuring the optical density in the presence (iron-limiting conditions) or absence (normal conditions) of 200 μM of the iron chelator 2,2′-bipyridyl (BIP) according to Conde-Pérez, Vázquez-Ucha [39]. Briefly, the bacteria were grown overnight in 5 mL LB broth at 37 °C with shaking at 200 rpm. The culture was then adjusted to an OD_600nm_ of 0.1 in Mueller-Hinton broth (MHB) and added to the wells with MHB, or MHB supplemented with 200 μM BIP (^-^Fe/^+^BIP), or 200 μM FeCl3 dissolved in 10 mM HCl (^+^Fe/^-^BIP), or 200 μM BIP plus 200 μM FeCl3 (^+^Fe/^+^BIP). The microplate was incubated at 37 °C in the Tecan Infinite 200 Pro, and the readings (OD_600nm_) were measured every 30 minutes for 21 hours. The growth curve was plotted using Prism 10 (Version 10.2.0). The maximum specific growth rate (μ_max_) and the lag time (λ) parameters were calculated using the single Gompertz growth curve model via the web tool (https://scott-h-saunders.shinyapps.io/gompertz_fitting_0v2). The maximum specific growth rate (μ_max_) represents the slope of the tangent at the inflexion point. The lag time (λ) represents the x-intercept of the μ_max_ tangent, which indicates the time (h) to enter the log phase. Three independent biological replicates were carried out.

### 2.10 Determination of total siderophore production by Chrome Azurol S assay

Chrome Azurol S (CAS) assays assessed the overall siderophore production in wild-type Ab-C98 and the knockout mutant derivatives using the microtiter plate method described in Senthilkumar, Amaresan [40]. Briefly, bacterial cultures well-grown in LB broth to stationary phase were adjusted to OD_600nm_ = 1.0, and 5 μL the culture was inoculated into 500 μL of MHB (normal condition) or MHB with 200 μM BIP (iron-limiting condition) and grown at 37 °C, 200 rpm for 21 hours. An inoculated broth control was included. The bacterial cultures were centrifuged at 10000 rpm for 10 mins, and the supernatant was collected. To perform the CAS assay, 100 μL of the supernatant was added to the microplate wells, and 100 μL of the CAS reagent (filter sterilised) was added into the wells. After 30 mins of incubation, the absorbance at 630 nm was taken (Tecan Infinite 200 Pro). The assay was performed in 5 independent biological replicates. The total siderophore was estimated by the following formula [40]:

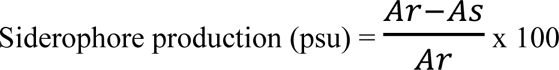

where Ar = absorbance of reference (CAS reagent and un-inoculated broth), and As = absorbance of sample (CAS reagent and cell-free supernatant of sample).

### 2.11 Prediction of the presence of *basC*, *bfnD* and NNO94_08395 isochorismate family protein in *Acinetobacter* species

The presence of the *basC*, *bfnD* and NNO94_08395 isochorismatase family protein was predicted in the pathogenic and non-pathogenic *Acinetobacter* species via Blastp, using the protein sequences of the three genes from Ab-C98 as query sequences and 90 % both for identity and query coverage as thresholds (e-value < 0.001). The complete genomes were retrieved from the NCBI database and annotated using Prokka (Galaxy Version 1.14.6+galaxy1) [41]. The complete genomes of the eleven *Acinetobacter* species were retrieved from the NCBI database. These include *A. baumannii* (710), *A. nosocomialis* (21), *A. pittii* (63), *A. lactucae* (2), *A. lwoffii* (11), *A. seifertii* (41), *A. junii* (15), *A. johnsonii* (26), *A. calcoaceticus* (7), *A. colistiniresistensis* (1) and *A. bouvettii* (1). The phylogenetic relationship between the representative strains of pathogenic and non-pathogenic *Acinetobacter* species was assessed. The gene content matrices were identified using Roary (Galaxy Version 3.13.0+galaxy2) [42]. The approximately-maximum-likelihood phylogenetic trees were constructed using FastTree (Galaxy Version 2.1.10+galaxy1) [43] and visualised using iTOL v6 [44]. The phylogenetic tree was rooted using the outgroup *Moraxella lacunata* strain NCTC 7911. The average nucleotide identity of the Ab-C98 and the reference strains of other *Acinetobacter* spp were calculated using the online web tool [45], and the heat map was generated using Prism 10 (Version 10.2.0). The genome alignment was performed using Mauve [46] using default parameters, and the presence of iron clusters was assessed.

### 2.12 Statistical analysis

Statistical analyses were performed using GraphPad Prism, version 10.2 (GraphPad Software Inc., San Diego, CA, USA). A correlation test was conducted using either Pearson or Spearman. Gene expression levels between infection and no infection control from RT-qPCR were analysed using an unpaired two-tailed Student’s t-test. Pearson correlation was used to analyse the correlation between direct RNA-seq and RT-qPCR. Growth kinetics between knockout mutants and the wild-type were analysed using one-way ANOVA followed by the Dunnett test. Analysis of survival curves was performed using a log-rank (Mantel-Cox) test. Bacterial burden data were first transformed by log_10_ and then analysed using Tukey’s multiple comparison test as part of one-way ANOVA to compare the differences between strains. A *p*-value < 0.05 indicates statistical significance. All values are expressed as means ± SEM.

## 3. Results

### 3.1 The reference clinical strain *A. baumannii* had lower killing than the Ab-C98

In our earlier report, Ab-C98 killed 83.33% of the larvae at an infection dose of 10^9^ CFU/mL after 24 hours of infection (*p* < 0.0001) [27]. Interestingly, lower infection doses of Ab-C98 were shown to be still infective to the larvae, where 10^8^ and 10^7^ CFU/mL inoculation of Ab-C98 resulted in 63% and 23% mortality after 3 days of infection (*p*<0.0001 and *p*=0.0005, respectively) [27]. To compare the killing of Ab-C98 with hospital strain *A. baumannii*, in this study, we performed a *G. mellonella* killing assay and bacterial burden assay for the clinical reference virulent strain ATCC BAA1605 (Figure 1). We found that at the same infection dose (10^9^ CFU/mL), ATCC BAA1605 displayed lower killing compared to the Ab-C98, with only 50 % of larvae killed after 24 hours of infection (**Figure 1**a). Lower infection doses of ATCC BAA1605 did not reduce the larval survival compared to the controls. To analyse the bacterial growth in the infected larvae, we performed statistical tests on our earlier reported results of Ab-C98 bacterial burden [27]. We found that at the infection dose of 10^9^ CFU/mL, there was a significantly increased Ab-C98 load at 3, 6 and 24 h p.i. compared to the 0 h control (*p*=0.0187, *p*=0.0375 and *p*=0.0011, respectively). In contrast, there is no significant difference in the ATCC BAA1605 bacterial load between time points (**Figure 1**b). Therefore, we aimed to identify the virulence factors of the Ab-C98 that caused the rapid killing and bacterial growth *in vivo* at the early infection stage (3 h p.i.) via transcriptomic analysis.

**Figure 1:**
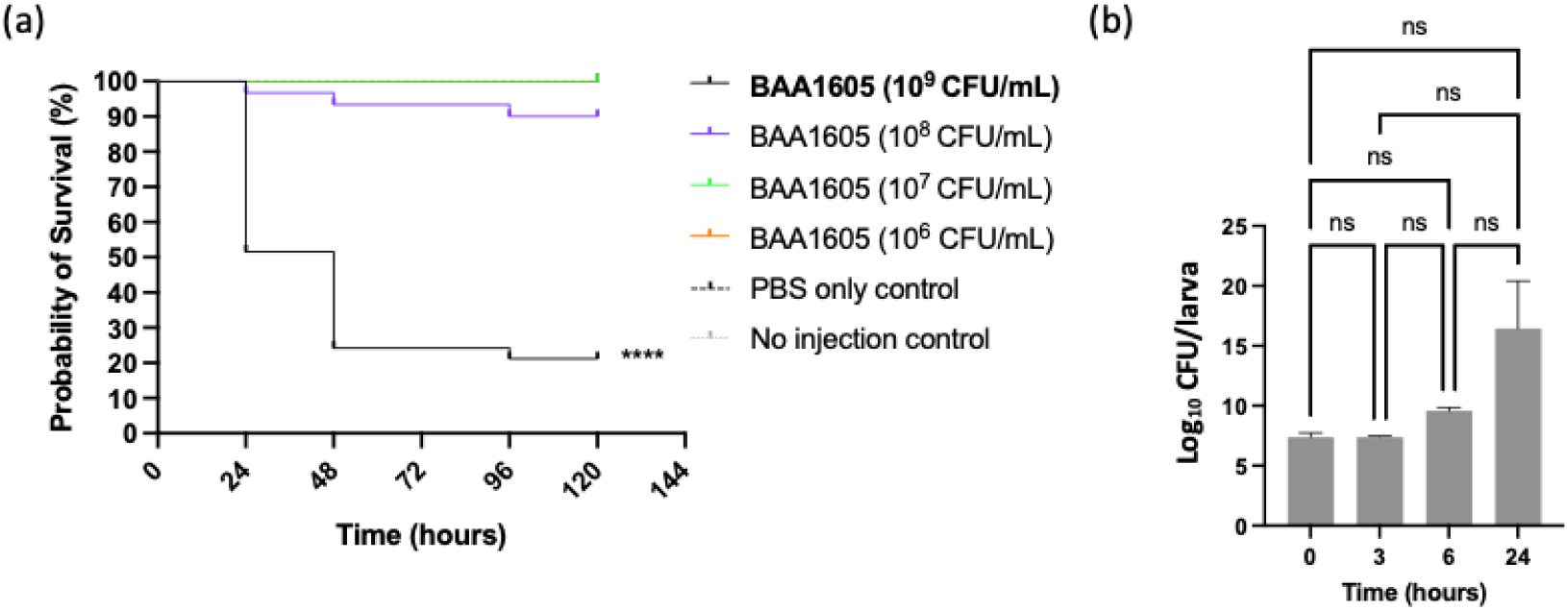
*G. mellonella* infection assays of *A. baumannii* clinical reference strain ATCC BAA1605. (a) Kaplan-Meier survival curves of *G. mellonella* infected by a serial dilution of ATCC BAA1605. The larval survival was scored every 24 hours for 5 days. Three independent replicates were performed (n=30). A log-rank test was performed to analyse the statistical significance of the larval survival compared to the controls upon 120 hours of infection. (b) The bacterial load (log_10_ CFU/larva) from the larvae infected by ATCC BAA1605 was quantified. Three independent replicates were carried out. The significant difference in bacterial load between the time points was analysed by one-way ANOVA.

### 3.2 Nanopore sequencing and mapping statistics of Ab-C98

To investigate the virulence mechanisms of Ab-C98, the transcriptome of this bacterium upon the infection was analysed by direct RNA-seq. The total RNA of Ab-C98 was collected from the infected larvae after 3 h p.i. (IVV), and the mid-exponential phase bacterial cells served as the no-infection control (IVB). The extracted RNA showed good purity with absorbance A260/230 and A260/280 = ∼2.0 (Supplementary Table S3). Each of the four libraries (infection samples replicate 1 and 2, IVV-1 and IVV-2, and no infection controls replicate 1 and 2, IVB-1 and IVB-2) was barcoded. The four libraries were sequenced on a single ONT flow cell and generated 2.37 million raw reads (BC1-4), with a mean read quality above 10 (Supplementary Table S4). The four libraries were evenly sequenced (Table 1), except the library BC4 (IVV-2, infection sample replicate no. 2) had a slightly lower read output. Generally, all libraries have a high mapping rate (>90%) to the Ab-C98 genome (Table 1). The *in vivo* libraries have negligible reads mapped to *G. mellonella*, except for BC4 (IVV-2), which had slightly higher host RNA contamination (88.16%, Table 1). Nevertheless, this data confirms the enrichment of *A. baumannii* bacterial cells from the infected host samples using our previously established protocol [27], with most reads mapped to the bacterium and minimal host RNA contamination.

**Table 1:** Mapping statistics of the Ab-C98 barcoded libraries after demultiplexing.

| Libraries | Raw reads | After trimming | Percentage of library in the pool (%) | Mapped reads to Ab-C98 | Mapped reads to Ab-C98 (%) | Mapped reads to *G. mellonella* | Mapped reads to *G. mellonella* (%) | Unmapped reads | Unmapped reads (%) |
| --- | --- | --- | --- | --- | --- | --- | --- | --- | --- |
| BC1 | 354810 | 353765 | 24.67 | 326487 | 92.29 | - | - | 27278 | 7.71 |
| BC2 | 409831 | 381881 | 26.63 | 352706 | 92.36 | 3894 | 1.02 | 25281 | 6.62 |
| BC3 | 428411 | 402131 | 28.04 | 378308 | 94.08 | - | - | 23823 | 5.92 |
| BC4 | 319691 | 296261 | 20.66 | 261185 | 88.16 | 13950 | 4.71 | 21126 | 7.13 |

Note: BC1: no infection control 1 (IVB-1); BC2: infection sample 1 (IVV-1); BC3: no infection control 2 (IVB-2); BC4: infection sample 2 (IVV-2).

### 3.3 Differential gene expression analysis demonstrates significant enrichment of iron acquisition systems in Ab-C98 during infection

The transcriptional responses of Ab-C98 during the infection were characterised via differential gene expression analysis to identify the genes that responded differently towards the stressful environment in *G. mellonella*. To analyse the correlation between the two biological replicates of each treatment group (IVV and IVB), the normalised read counts of the transcripts were obtained from the featureCount, and a Spearman correlation test was performed. There is a strong correlation between the two biological replicates of the no infection control (Spearman’s correlation = 0.8328, *p* < 0.0001) and the infection samples (Spearman’s correlation = 0.7097, *p* < 0.0001) (**Figure 2**a), suggesting that the two biological replicates had similar expression profiles. Fifty-three significant DEGs were identified, with 45 genes up-regulated and 8 genes down-regulated in the infection conditions compared to the no infection control (**Figure 2**b). Of the up-regulated genes, 29 belong to the iron acquisition system, with genes involved in the acinetobactin synthesis (*bas*, *bau*, *bar* genes) as the major differentially expressed iron cluster upon the infection (**Figure 2**c). Additionally, we recorded a significant up-regulation of the osmotic stress response operon (*betIBA*) (log_2_fold-change=2.13-2.39) and BCCT family transporter (NNO94_03680) (log_2_fold-change=3.04), and a significant down-regulation of OmpW family outer membrane protein (NNO94_00550) (log_2_fold-change= -2.91) (**Figure 2**c). Functional enrichment analysis also showed that the iron transport and siderophore production pathways were significantly enriched in the infection samples compared to the no-infection controls (**Figure 2**d). Sixteen of the 18 genes within the acinetobactin cluster were up-regulated by at least 7.26-fold (**Figure 2**e, up-regulated genes were coloured in red). These include the *bas* genes (*basB,C,E*,*F*,*G*,*D*,*J*,*H,I*) that are involved in the pre-acinetobacter synthesis and acinetobactin assembly, the *barAB* genes that are involved in the siderophore efflux system, and the *bau* genes (*bauA*,*B*,*C*,*D*,*E*) that are responsible for the recognition and translocation of ferric acinetobactin [39,47,48]. Besides, we observed a second cluster of genes (located within 1639507 bp to 1654730 bp of the Ab-C98 chromosome) up-regulated in Ab-C98 during the infection. This cluster has a high Blastp similarity (>90%) to the baumannoferrin cluster (*bfnA*-*L*) in *A. baumannii* ACICU (**Figure 2**e). The six up-regulated baumannoferrin genes from this cluster include NNO94_07825 siderophore biosynthesis protein (*bfnA*), NNO94_07830 SidA/IucD/PvdA family monooxygenase (*bfnB*), NNO94_07840 siderophore achromobactin biosynthesis protein AcsC (*bfnD*), NNO94_07845 IucA/IucC family siderophore biosynthesis protein (*bfnE*), NNO94_07850 (2Fe-2S)-binding protein (*bfnF*) and NNO94_07880 acetyltransferase (*bfnL*).

**Figure 2:**
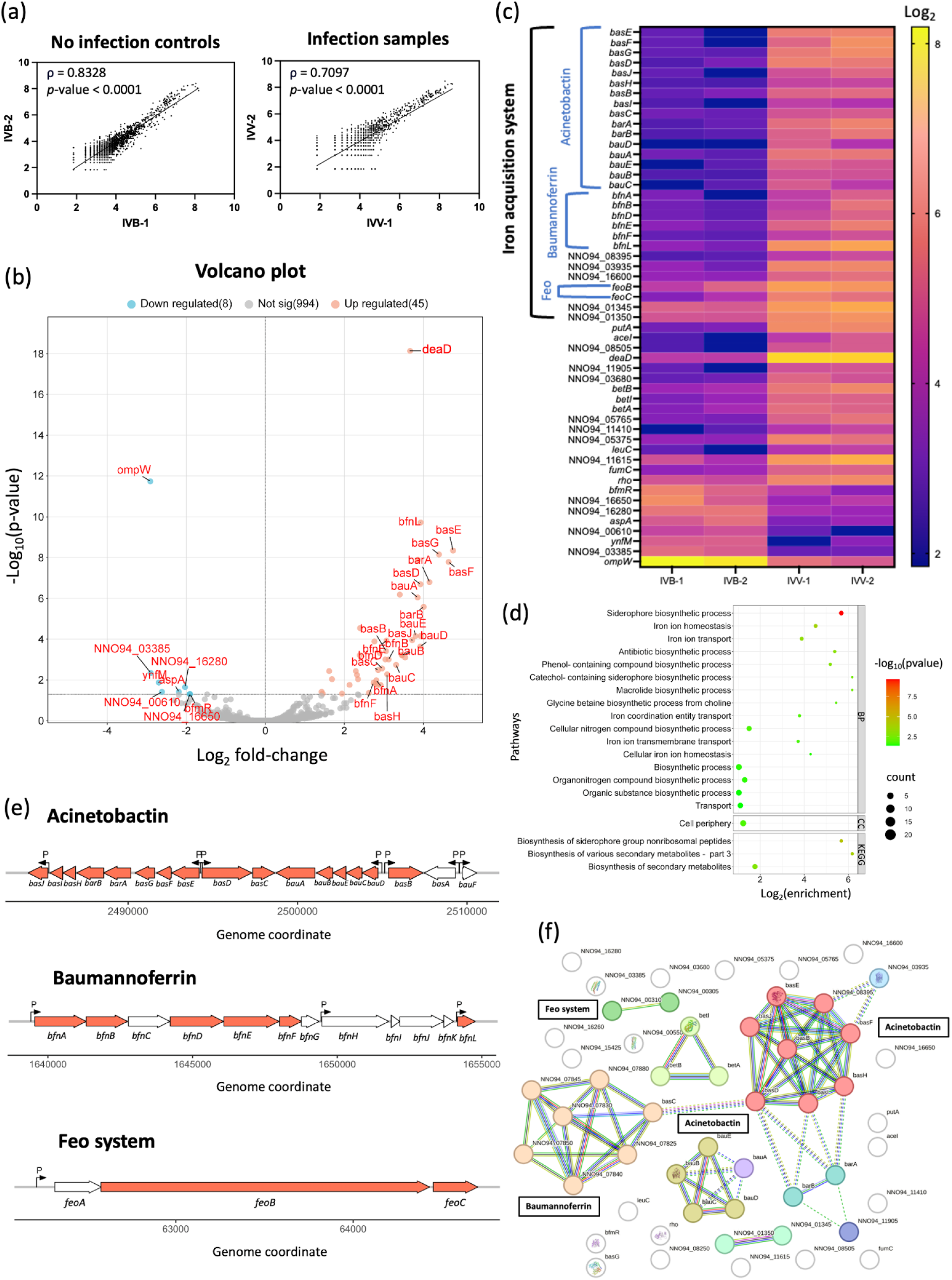
Differential gene expression analysis revealed the significant up-regulation of three iron clusters in Ab-C98 during infection. (a) Correlation between the two biological replicates of the no infection controls (IVB) and infection samples (IVV). (b) Volcano plot of differential gene expression analysis. Red = up-regulated genes; Blue = down-regulated genes; Grey = not significantly differentially expressed. (c) Heatmap of differentially expressed genes, plotted using VST transformed normalised read counts in log_2_-scaled. The higher the normalised read counts, the higher the expression of the genes. (d) Functional enrichment analysis of the 53 differentially expressed genes in Ab-C98 upon infection. BP = Gene Ontology Biological process, CC = Gene Ontology Cellular Components, KEGG = Kyoto Encyclopedia of Genes and Genomes. (e) Genome coordination of the three iron clusters (acinetobactin, baumannoferrin and Feo system) in the Ab-C98 chromosome, with genes differentially expressed during infection coloured in red. Non-differentially expressed genes are coloured in white. The predicted promoter regions were annotated as ‘P’. (f) Functional association among the 53 differentially expressed genes in Ab-C98 upon infection. k-means clustering was enabled to cluster interacting proteins in different groups. Dashed lines represent inter-cluster edges.

An increased expression of genes involved in the transportation of iron was also observed. A significant up-regulation of *feoB* (NNO94_00305 ferrous iron transporter B) and *feoC* (NNO94_00310 hypothetical protein, homolog of DMO12_RS01465 in ACICU strain) was noticed in Ab-C98 upon the infection (log_2_ fold change of 1.69 and 2.29, respectively) (**Figure 2**e). *feoA* was not found to be significantly differentially expressed in Ab-C98. Furthermore, two genes responsible for the transport of iron-siderophore complex through the outer membrane into the bacterial cell [49], NNO94_01350 biopolymer transporter ExbD and NNO94_01345 MotA/TolQ/ExbB proton channel family protein were up-regulated in Ab-C98 upon the infection. Functional protein association network analysis showed that the enriched DEGs form four intermediated clusters of iron acquisition systems (**Figure 2**f).

To independently validate the expression levels of the DEGs observed in the transcriptomic analysis, we investigated the expression levels of selected targets from the two major siderophore clusters (acinetobactin and baumannoferrin) via RT-qPCR. These target genes include the *basE* and *basC* (acinetobactin cluster) and the *bfnA* and *bfnD* (baumannoferrin). The RT-qPCR analysis showed that *basE*, *basC* and *bfnD* were significantly up-regulated in the infection sample compared to the no-infection control (log_2_ fold change of 9.11, 8.37 and 4.66, respectively). In contrast, bfnA had an insignificant change in expression level upon infection (*p*=0.4) (**Figure 3**a, Supplementary Figure S1). In addition, we selected a gene encoding the isochorismatase family protein (NNO94_08395), which does not belong to any known iron clusters in *A. baumannii*. We confirmed its significant up-regulation by the RT-qPCR (log_2_ fold change of 5.60) (**Figure 3**a, Supplementary Figure S1). We also validated the expression levels of two targets unrelated to the iron clusters, which are the *deaD* (NNO95_15425, responsible for the RNA metabolism), represent the most significant up-regulated DEG (*p*=7.39 × 10^-19^) and *ompW* (NNO94_00550, an outer membrane beta-barrel protein) represent the most significant down-regulated DEG (*p*=1.84 × 10^-12^) from the DGE analysis. These two genes were confirmed to be significantly differentially expressed upon infection from the RT-qPCR, with a log_2_ fold change of 8.00 and -6.69, respectively (Figure 3a, Supplementary Figure S1). A high correlation between the direct RNA-seq and RT-qPCR results was found using Pearson correlation (r=0.9466, *p*-value=0.0012) (**Figure 3**b).

**Figure 3:**
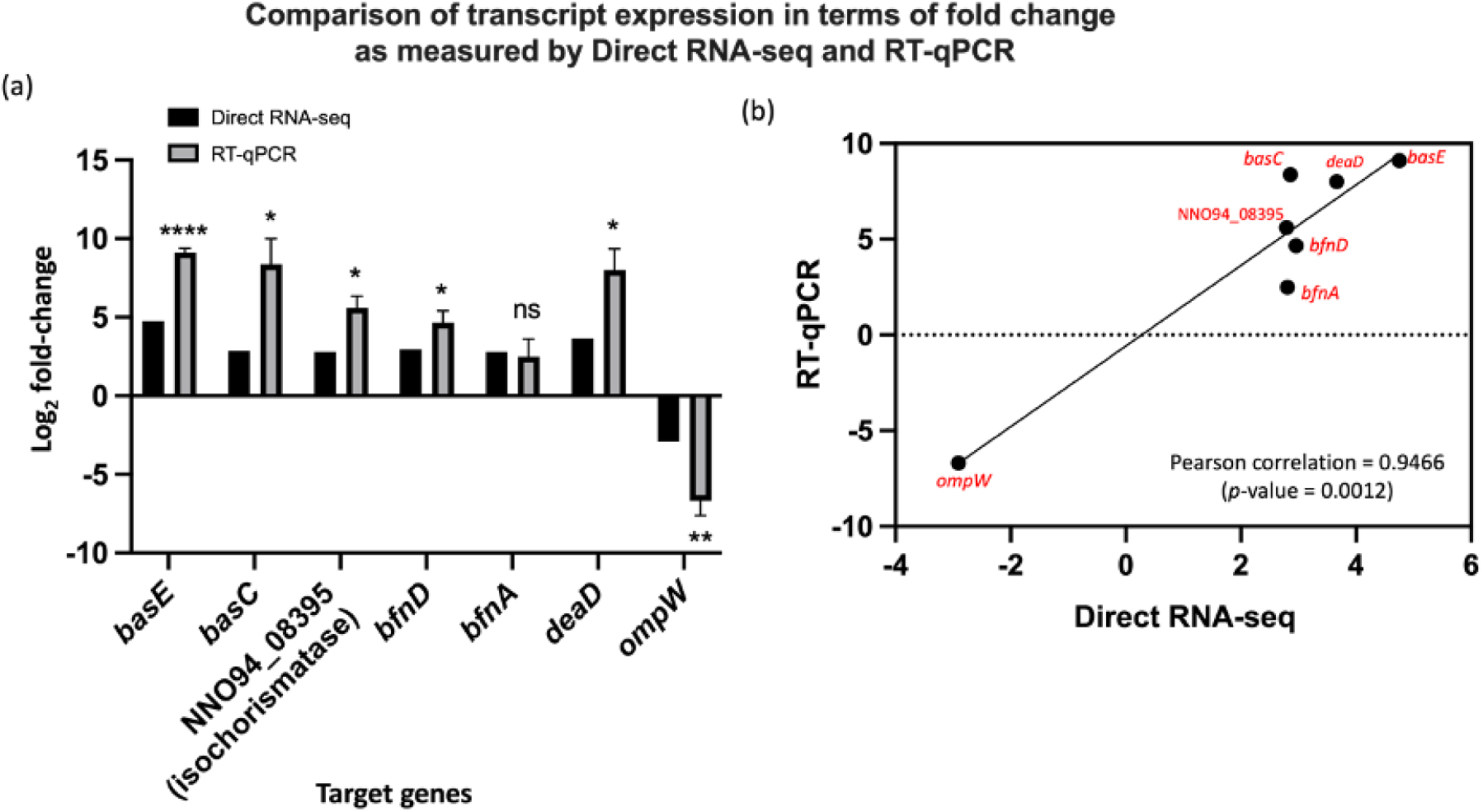
RT-qPCR and dRNA-seq fold change values are highly correlated. The expression levels of selected targets in Ab-C98 upon the infection in *G. mellonella* were analysed via RT-qPCR. Bar plots and inset scatter plots compared dRNA-seq and RT-qPCR fold change values obtained for the seven shortlisted differentially expressed genes in log_2_-scaled. Three independent biological replicates with three technical replicates each were performed. The expression levels of the genes between the infection samples and no infection controls were analysed by unpaired, two-tailed Student-t tests. Asterisks on the qPCR values indicate a significant difference between the infection sample and non-infection control Ab-C98. p<0.05 (*), p<0.01 (**), p<0.001 (***), p<0.0001 (****), ns (non-significant).

### 3.4 BasC is essential for *A. baumannii* growth under iron-depleted conditions *in vitro*

Iron acquisition represented the critical pathway in Ab-C98 during the infection, so we further selected one representative gene from each cluster for functional assessment via gene knockout: (i) *basC*, responsible for the biosynthesis of acinetobactin precursor; (ii) *bfnD*, responsible for the biosynthesis of baumannoferrin, and (iii) NNO94_08395 isochorismatase family protein. Interestingly, the role of the *bfnD* and isochorismatase in *A. baumannii* virulence has not yet been studied. Isogenic knockout mutants were obtained for each of the targeted genes.

To determine the effect of the knockout genes on the growth of Ab-C98, the growth curves of the wild-type Ab-C98 and the three knockout mutants (Δ*basC*, Δiso, Δ*bfnD*) were performed in the absence (normal conditions) and the presence (iron-limiting condition) of the iron chelator 2,2′-bipyridyl (BIP) (**Figure 4**). Under normal conditions (MHB broth medium), wild-type Ab-C98 and the three mutant strains had similar growth rates (μ_max_) and lag time (λ), indicating that deletion of the genes did not affect bacterial standard growth *in vitro* (**Figure 4**a, Supplementary Figure S2a-b). However, the Δ*basC* strain lacking the gene involved in the synthesis of the N-hydroxyhistamine precursor necessary for the acinetobactin assembly showed a significantly lower growth rate (μ_max_ = 0.06, *p*<0.0001) than the wild-type strain (μ_max_ = 0.13) under the iron-limiting conditions (**Figure 4**b, Supplementary Figure S2c). Interestingly, Δiso (NNO94_08395 isochorismatase family protein) and Δ*bfnD* mutant strains displayed similar growth rates (μ_max_ = 0.1460 and 0.1534, respectively) to the wild-type strain under iron-limiting conditions (**Figure 4**b, Supplementary Figure S2c). All strains had a similar lag time as the wild type in the iron-limiting conditions (Supplementary Figure S2d). No significant differences in growth kinetics were observed in all strains when iron source (FeCl_3_) was added [p(μ_max_) = 0.13-0.99, p(λ) = 0.67-0.93] (**Figure 4**c and **Figure 4**d, Supplementary Figure S2e-h).

**Figure 4:**
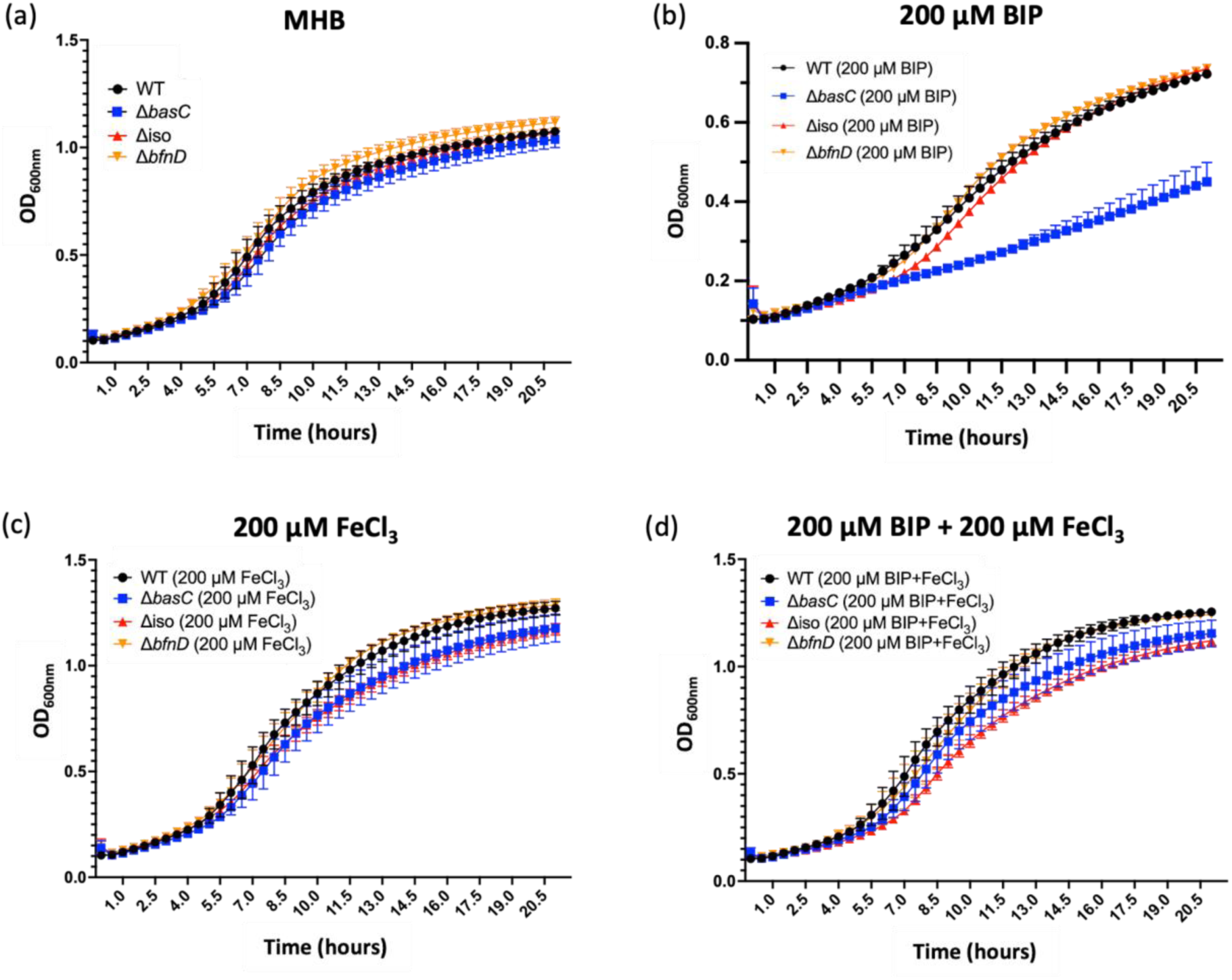
Growth curves of Ab-C98 and its isogenic mutant derivatives under the conditions of (a) MHB culture medium only, (b) 200 µM BIP (iron-depleted), (c) 200 µM FeCl_3_ (iron-rich), and (d) 200 µM BIP and 200 µM FeCl_3_. Three independent biological replicates were carried out. The error bars represent the mean ± SEM.

To evaluate the changes in the siderophore production upon the deletion of the genes, we performed the CAS microplate method to assess the overall production of siderophores in wild-type Ab-C98 and its knockout mutant derivatives under normal conditions (MHB) and iron-limiting conditions (MHB supplemented with 200 μM BIP) (Supplementary Figure S3). All strains demonstrated a significant increase in the total siderophore activity under iron-depleted conditions (*p*=0.0121-0.0269); however, insignificant differences in total siderophore activity were observed between wild-type and knockout mutant strains under normal and iron-depleted conditions.

### 3.5 *basC*, isochorismatase family protein and *bfnD* are required for *A. baumannii* virulence during infection

Acinetobactin has been reported to play significant virulence roles in *A. baumannii* clinical reference strain in murine models [39]. However, whether these genes have similar effects in a community *A. baumannii* strain and a non-mammalian model remains unclear. To assess the virulence of the three siderophore production genes in the wild-type Ab-C98 and the three knockout mutants, *G. mellonella* infection assays were performed at the same infection dose for the four bacterial strains. Among the three mutant strains, the larvae infected with Δ*basC* mutant had the most significantly reduced killing (76.67% survival, *p* < 0.0001) compared to the wild-type strain (10% survival) after 6 h of infection (**Figure 5**a). This is followed by the Δiso (53.33% survival, *p*=0.0002) and Δ*bfnD* (45.16% survival, *p*=0.0018). To elucidate whether the reduction in the killing is due to a change in the bacterial load, a *G. mellonella* bacterial burden assay was performed to quantify the CFU/larva at 0, 3 and 6 h post-infection (p.i.) (**Figure 5**b). There were no significant differences in the bacterial load in all the three mutant strains and the wild-type strain at the 0 h and 3 h p.i. (*p*=0.8733-0.9997). Although the three mutant strains displayed a slightly reduced bacterial load compared to the wild-type at 3 h p.i., the differences were insignificant. Similarly, a marginally lower bacterial load was observed in the mutant strains at 6 h p.i., compared to the wild-type (Δ*basC p*=0.0121, Δiso *p*=0.0255 and Δ*bfnD p*=0.0637).

**Figure 5:**
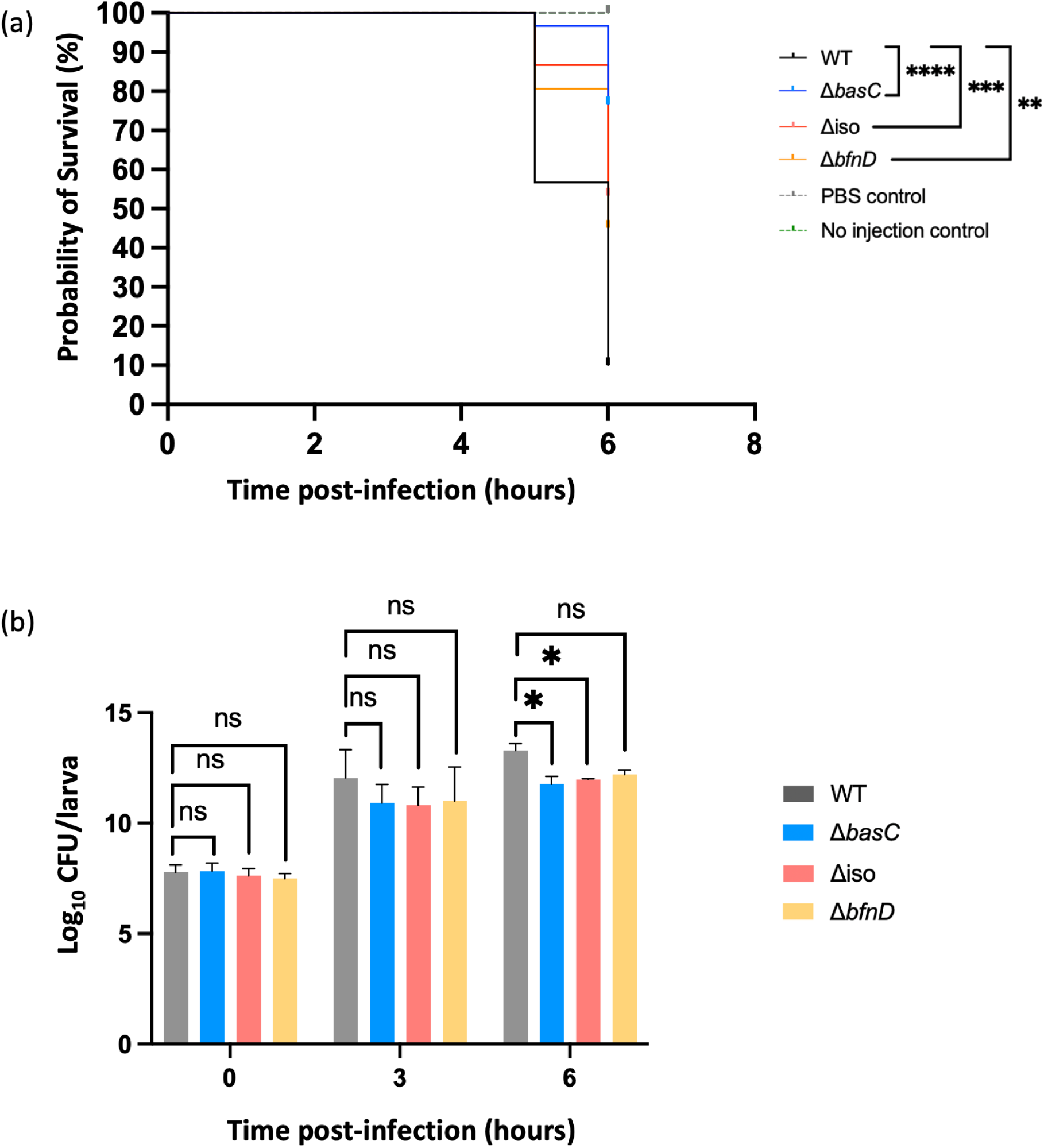
Virulence assessment of wild-type Ab-C98 and its mutant derivatives via *G. mellonella* infection assays. (a) Kaplan-Meier survival curves of *G. mellonella* infected by Ab-C98 and its mutant derivatives. A log-rank test was performed upon 6-h infection to analyse the statistical difference in *G. mellonella* survival between wild-type and the knockout mutants. Three independent replicates were carried out (n=30). (b) Bacterial load of Ab-C98 and its mutant derivatives at designated time points (0, 3, and 6h). Three independent replicates were performed. The significant difference in bacterial load between the wild-type and the knockout mutant strains was analysed by one-way ANOVA. Asterisks indicate a statistical significance between the knockout mutants and the wild-type. p<0.05 (*), p<0.01 (**), p<0.001 (***), p<0.0001 (****), ns (non-significant).

### 3.6 The three siderophore production genes are commonly present in *A. baumannii*

The virulence of the targeted siderophore production genes (*basC*, isochorismatase and *bfnD*) has been confirmed as described above. Therefore, we aim to assess if these identified virulence genes are also present in other *Acinetobacter* species to understand their broader implications for pathogenesis. To identify the presence of the genes, we performed Blastp, using the gene sequences from the Ab-C98 as the query sequences, against eleven *Acinetobacter* spp (**Table 2**). The phylogenetic tree of the eleven *Acinetobacter* spp was also constructed from the core gene analysis (**Figure 6**). The eleven *Acinetobacter* spp include the *Acinetobacter calcoaceticus*-*baumannii* complex (Acb complex), which consists of six members: *A. calcoaceticus*, *A. baumannii*, *A. nosocomialis*, *A. pittii* and *A. lactucae* (formerly known as *A. dijkshoorniae*), with *A. baumannii*, *A. nosocomialis* and *A. pittii* represent the most critical nosocomial pathogens [1,50]. Additionally, *A. lwoffii*, *A. seifertii*, *A. junii* and *A. johnsonii* are included in this analysis and categorised as pathogenic *Acinetobacter* due to their occasional reports of infections [51]. The environmental-origin *A. calcoaceticus* was used in this analysis as a non-pathogenic *Acinetobacter* due to its clinical unimportance [52,53]. Other rare or non-pathogenic *Acinetobacter* spp included in this analysis are *A. colistiniresistensis* and *A. bouvettii*. Of the eleven *Acinetobacter* spp, the baumannoferrin *bfnD* is present only in the *A. baumannii* strains (96.90% of the 710 complete genomes) and absent in the other *Acinetobacter* spp (**Table 2**). Interestingly, *basC* and the NNO94_08395 isochorismatase family protein are present not only in the *A. baumannii* but also in two other pathogenic *Acinetobacter* spp, which are the *A. pittii* (90.48% and 92.06% respectively), and the *A. lactucae* (100% and 100% respectively). The phylogenetic analysis showed that the *A. nosocomialis* strains are clustered within *A. baumannii* strains, suggesting they have a close phylogenetic relationship (**Figure 6**). However, the three siderophore production genes are absent in all the *A. nosocomialis* strains (Table 2, **Figure 6**). The non-pathogenic *A. calcoaceticus* is clustered in the same clade as the pathogenic *A. pittii* and *A. lactucae* and is the sister group with the *A. baumannii* cluster (**Figure 6**). Still, the three siderophore production genes are absent in the *A. calcoaceticus*. Furthermore, the genes are also not detected in the *Acinetobacter* spp in another cluster that contains *A. junii*, *A. colistiniresistensis*, *A. johnsonii*, *A., bouvettii* and *A. lwoffii* (**Figure 6**). This suggests that the three siderophore production genes are mainly present in the pathogenic *Acinetobacter* spp (*A. baumannii*, *A. pittii* and *A. lactucae*), and baumannoferrin *bfnD* is present only in the *A. baumannii* strains.

**Figure 6:**
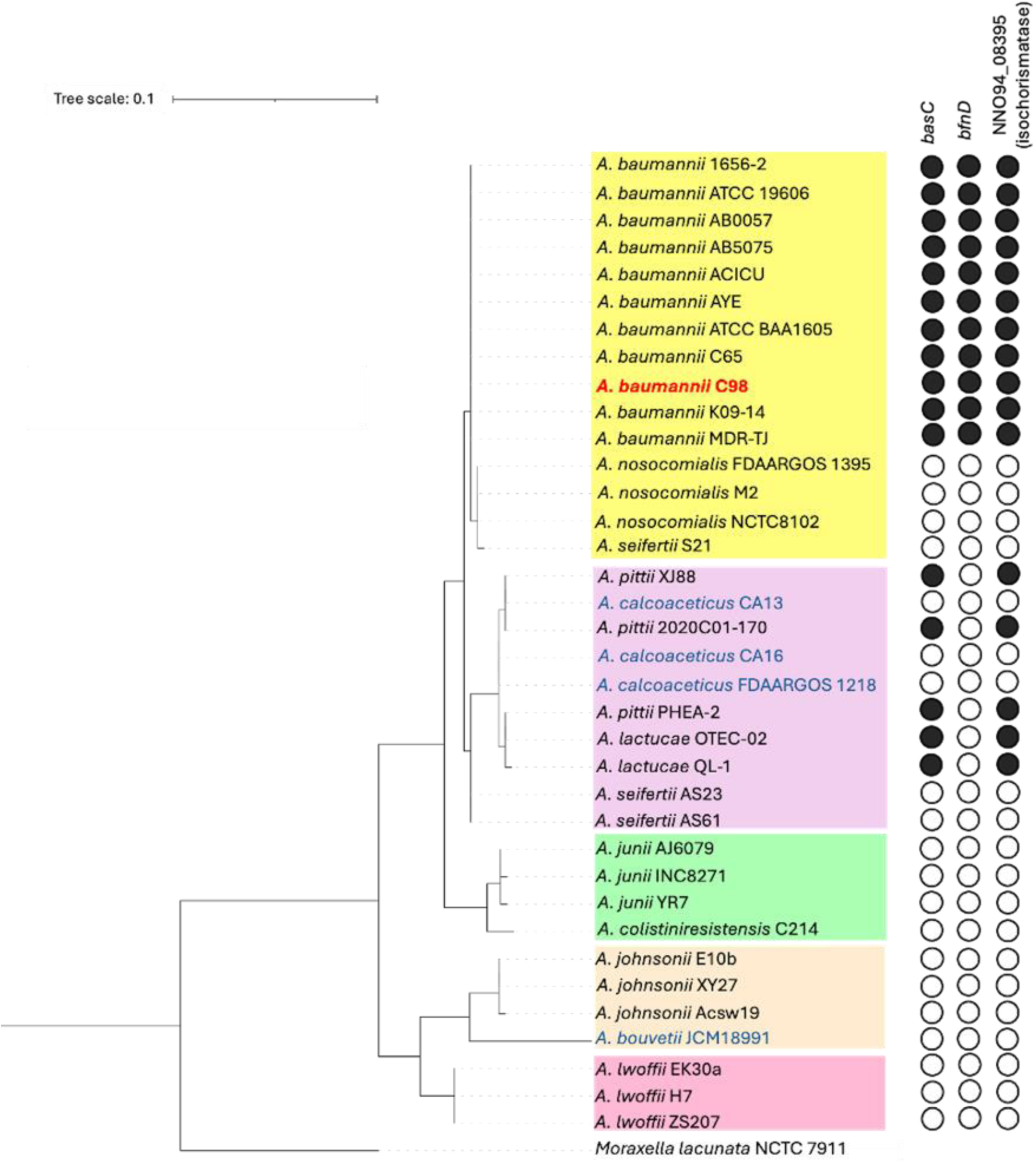
Phylogenetic analysis of *basC*, *bfnD* and isochorismatase in the 36 pathogenic and non-pathogenic *Acinetobacter* strains. Phylogenetic tree of pathogenic and non-pathogenic *Acinetobacter* spp and the absence/presence of the 3 siderophore production genes among the strains. For *A. baumannii*, Ab-C98 (highlighted in red) and 11 complete genomes were selected for the phylogenetic analysis. For other *Acinetobacter* species, 3 complete genomes were selected based on their respective phylogenetic clusters. Non-pathogenic *Acinetobacter* spp. were highlighted in blue. *Moraxella lacunata* strain NCTC 7911 was used as an outgroup. The phylogenetic tree was constructed based on the core genes analysed by Roary. The strains clustered in the same clade were coloured. The absence or presence of the genes was analysed using blastp, with query coverage and identity ≥ 90% (e-value < 0.001). Full circles indicate the presence of the gene, while empty circles indicate the absence of the gene.

**Table 2:**
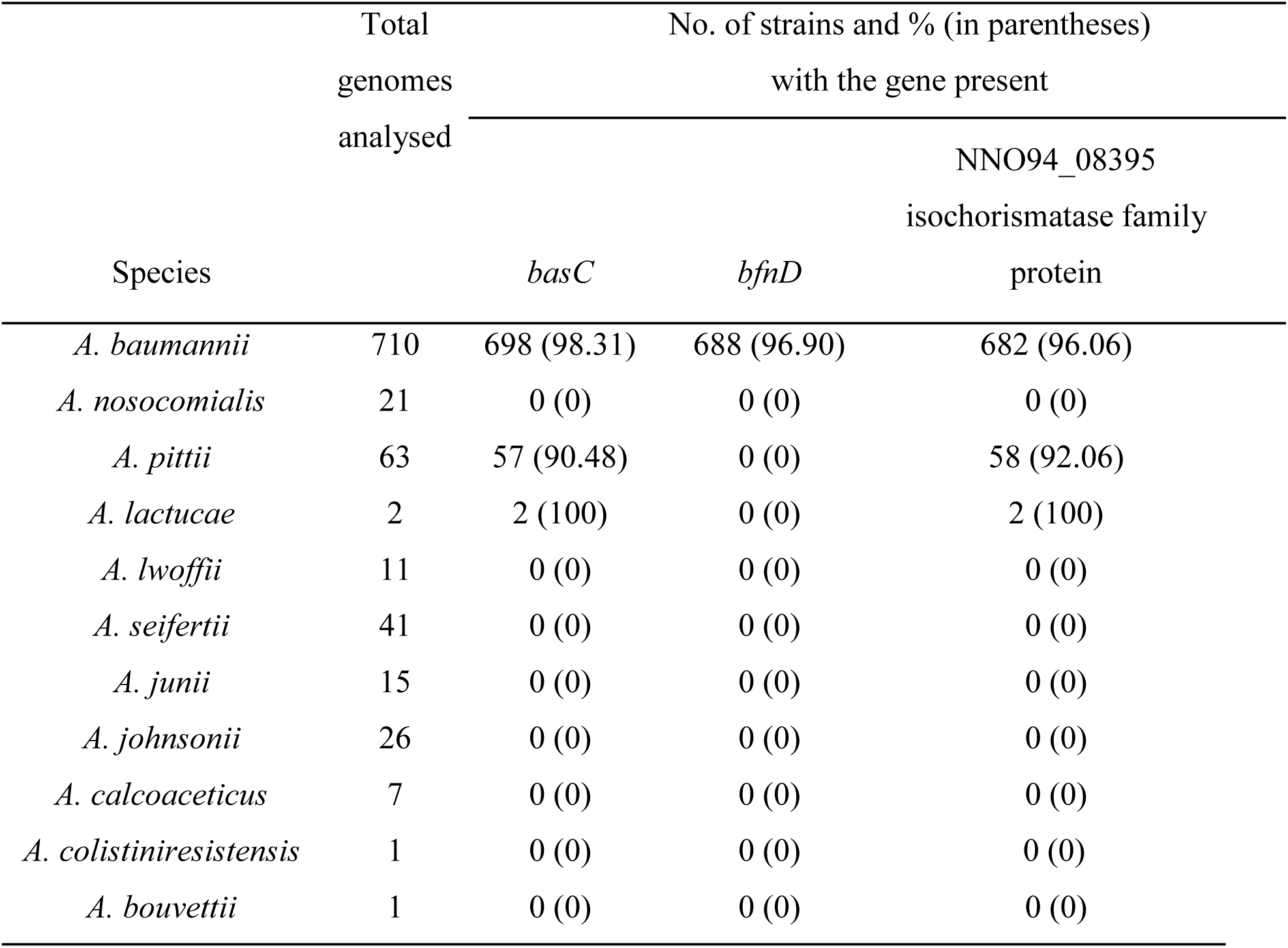
Prediction of the 3 siderophore production genes in pathogenic and non-pathogenic *Acinetobacter* species using blastp.

The average nucleotide identity analysis on the Ab-C98 and other *Acinetobacter* spp showed that the *A. nosocomialis* strain M2 has the highest genome similarity with Ab-C98, followed by the *A. seifertii* strain S21, *A. pittii* strain PHEA-2, *A. lactucae* strain QL-1 and *A. calcoaceticus* strain CA16 (ANI value >80%) (**Figure 7**a). This is consistent with our phylogenetic analysis using the core genes, indicating these five species are closely related. Other non-*A. baumannii* spp (*A. junii*, *A. colistiniresistensis*, *A. johnsonii*, *A. lwoffii* and *A. bouvetii*) have an ANI value of 73.39-76.69% (**Figure 7**a), indicates that they have distinct genomic profiles with Ab-C98. The presence of the acinetobactin and baumannoferrin clusters was assessed by aligning the whole genomes of *A. baumannii* and the *Acinetobacter* spp from its sister group (*A. nosocomialis*, *A. seifertii*, *A. pittii*, *A. lactucae* and *A. calcoaceticus*) using the reference strains. The acinetobactin cluster is conserved in the *A. baumannii* and the closely related species *A. pittii* and *A. lactucae*, as indicated by the high similarity profile in the acinetobactin locus (**Figure 7**b). Interestingly, despite having a close phylogeny with *A. baumannii*, the acinetobactin locus is completely absent in the *A. nosocomialis*. Besides, the acinetobactin locus is not found in the pathogenic *A. seifertii* and non-pathogenic *A. calcoaceticus*. Similarly, the baumannoferrin locus is only present in the *A. baumannii*, while absent in other *Acinetobacter* spp (**Figure 7**c).

**Figure 7:**
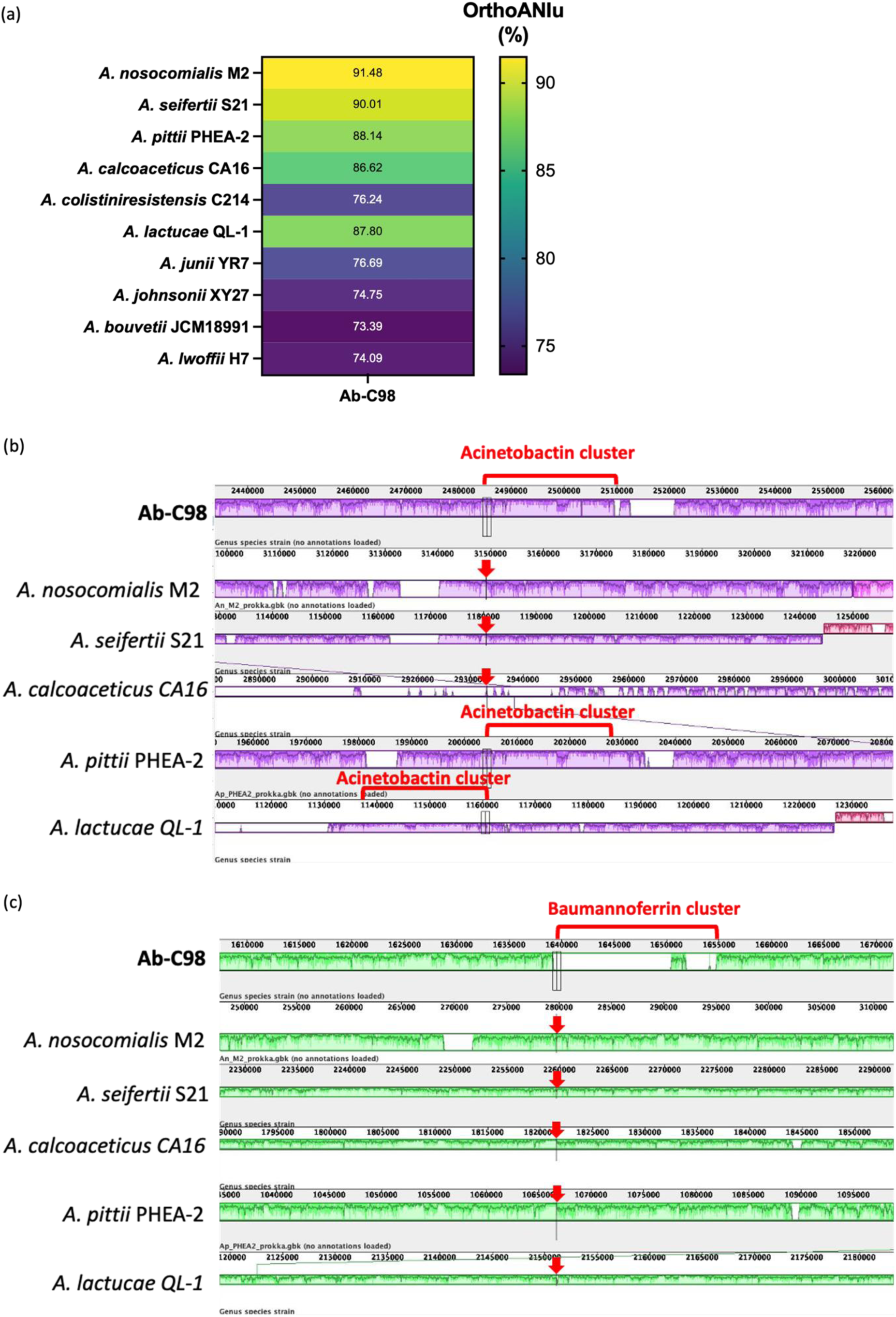
Genome alignment of the acinetobactin and baumannoferrin clusters in Ab-C98 and other *Acinetobacter* spp from its sister group using Mauve. (a) The average nucleotide identity of the Ab-C98 with the reference strains of other *Acinetobacter* spp. Genome alignment of the (b) acinetobactin cluster and (c) baumannoferrin cluster was visualised using Mauve. The presence of the iron clusters was detected by hovering the mouse cursor over the gene locus of the iron clusters, which was highlighted in the red brackets. A red arrow indicated the absence of the cluster. The reference strains of the pathogenic *A. nosocomialis* (strain M2), *A. seifertii* (strain S21), *A. pittii* (strain PHEA-2) and *A. lactucae* (strain QL-1), and the non-pathogenic *A. calcoaceticus* (strain CA16) with close phylogeny with *A. baumannii* were compared. The coloured blocks outline the regions that align to part of other genomes, and coloured lines connect the homologous regions in each genome. The height of the similarity profiles in each coloured block corresponds to the average level of conservation in that region of the genome sequence. A completely white region within the coloured block indicates a lack of alignment and is unique in a particular genome.

## 4. Discussion

### 4.1 Community-acquired *A. baumannii* showed higher virulence than the clinical strain in *G. mellonella* model

Community-acquired *A. baumannii* infections are usually uncommon, but they can be highly fatal, with mortality rates of 60% and above [12,54]. We previously established the *A. baumannii*-*G. mellonella* infection model using the Ab-C98 as the pathogen and found that this Ab-C98 could kill the larvae in a dose-dependent manner [27]. This strain has a close phylogenetic relationship with a cluster of hospital-acquired *A. baumannii* isolated from the local hospital. It possesses a complete set of type VI secretion systems absent in the other community strains [55]. We then evaluated the killing of the clinical reference virulent strain ATCC BAA1605 in *G. mellonella* and compared it to Ab-C98. Interestingly, the clinical strain ATCC BAA1605 displayed a lower killing than the Ab-C98 and had insignificant differences in bacterial load in the infected larvae after 24 hours of infection. The rapid colonisation of the Ab-C98 in the larvae may contribute to the higher killing of this strain. Similarly, Peleg and Jara [56] reported a greater killing of community-acquired *A. baumannii* than the hospital strain in the *G. mellonella* model. In contrast, Chusri observed a similar killing between community- and hospital-acquired *A. baumannii*. These findings suggest that the community-acquired *A. baumannii* had virulence similar to hospital strains despite their high antibiotic susceptibility. Therefore, it is important to understand the virulence mechanisms of community-acquired *A. baumannii*.

### 4.2 Transcriptomic analysis and functional validation reveal the *basC*, isochorismatase and *bfnD* as virulence factors in Ab-C98 pathogenesis in *G. mellonella* model

Transcriptomic analysis allows us to analyse the genes differentially regulated by *A. baumannii* during an infection. The high correlation with the independent RT-qPCR validations has indicated the reliability of the gene expression analysis from the nanopore sequencing data. We observed a significant enrichment in the iron acquisition system in the Ab-C98 upon the infection in *G. mellonella*. This includes the two major siderophore clusters, acinetobactin and baumannoferrin, which are significantly up-regulated, and one iron transport system (Feo system). Our findings suggest that the low iron availability in the host environment (*G. mellonella* larvae) upon the *A. baumannii* infection drives the bacterium to uptake additional extracellular iron via increased siderophore production and iron transportation, in agreement with Dunphy, Niven [57] where *Xenorhabdas nematophila* pathogenic infection caused a reduced iron content in the larval hemolymph; however, the overall bacterial development was not affected due to the production of siderophores [57]. Furthermore, our findings align with Sheldon and Skaar [58], where up-regulation of acinetobactin and baumannoferrin genes were observed in *A. baumannii* from infected mice via NanoString; however, only a few genes from the two clusters were tested. Our transcriptomic analysis further confirmed the significance of the iron clusters in *A. baumannii* virulence and survival in infection by demonstrating the significant up-regulation of nearly the entire cluster via direct RNA-seq, using the non-mammalian insect model *G. mellonella*.

From the protein-protein interaction analysis, we observed that the genes involved in the acinetobactin and baumannoferrin clusters interact, with *basC* and *basD* in the centre connecting the two groups. This indicates the two clusters contribute jointly to the same biological function, which is the siderophore production that enhances the bacterial survival upon the infection. Additionally, a gene encoding the isochorismatase family protein (NNO94_08395) was clustered within the acinetobactin *bas* genes. This gene has a high Blastp identity (99.061%) with the DMO12_RS09225 isochorismatase family protein in *A. baumannii* strain ACICU, which does not belong to the known iron clusters [59]. It has been reported that deleting genes involved in siderophore production reduced the virulence of pathogens such as *Staphylococcus* and *Salmonella* [60]. This has also been shown in Conde-Pérez and Vázquez-Ucha [39], where deletion of genes from the acinetobactin cluster (*basG*, *basC*, *basD*, *basB*, *bauA*) significantly improved the survival of *A. baumannii*-infected mice. At the same time, growth inhibition *in vivo* was observed upon deletion of *basG* [58]. However, there are no previous reports on the role of the isochorismatase and baumannoferrin genes in *A. baumannii* virulence.

We investigated the knockout mutants’ growth upon gene deletion from the acinetobactin cluster (*basC*), baumannoferrin cluster (*bfnD*) and the isochorismatase family protein *in vitro* and *in vivo*. We observed a reduced bacterial growth rate *in vitro* upon deletion of *basC* in *A. baumannii* under iron-depleted conditions. The *basC* gene synthesises N-hydroxyhistamine, a precursor required to assemble acinetobactin [39]. The absence of *basC* inhibited the acinetobactin synthesis. However, fimbactins A and F were still detected in the *A. baumannii* Δ*basC* knockout mutants [39]. This could explain the insignificant differences in our findings on the total siderophore activity between the Δ*basC* and the wild-type strain. The small changes in the siderophore production by a single deletion of the gene involved in the particular siderophore may not be effectively detected using the CAS assay. Ab-C98 Δ*bfnD* and Δiso (isochorismatase family protein) mutants grew well in the iron-depleted medium *in vitro*. Similar observations were reported in Sykes [22], where they observed higher growth inhibition in the mutants lacking the acinetobactin than the fimsbactin and baumannoferrin mutants under the metal-chelated minimal medium with human serum. Our findings indicate that acinetobactin was the primary siderophore utilised by the Ab-C98 under iron-depleted conditions. Our G. mellonella infection further observes these assays on the wild-type Ab-C98 and the three isogenic mutant derivatives, where the wild-type strain had the highest killing rate, followed by the Δ*basC*, Δiso and Δ*bfnD*. Interestingly, the bacterial load in the three knockout mutants was marginally lower than that of the wild-type strain, which indicates that the absence of the genes has minimal impact on bacterial growth *in vivo* but can reduce bacterial virulence by reducing the killing rate in the larvae at the early infection stage. This could lower the possibility of bacterial resistance development against anti-virulence agents targeting these siderophore production genes due to weaker selective pressure [61].

### 4.3 High conservation of acinetobactin, baumannoferrin and isochorismatase as a contributor to *A. baumannii* virulence, but not observed in closely related pathogen *A. nosocomialis*

*Acinetobacter* species usually exist in the natural environment, such as soils, oceans and freshwater [52]. However, due to its various innate mechanisms, *Acinetobacter* spp has evolved as an emerging pathogen capable of causing diseases, with *A. baumannii* having the most clinical significance, which accounts for 80% of hospital-acquired infections [62]. The observed virulence of the selected siderophore production genes prompts whether the identified virulence genes are conserved across other *Acinetobacter* species capable of causing diseases. The Blastp analysis showed that *basC* and isochorismatase are highly present in three pathogenic *Acinetobacter* spp. (*A. baumannii*, *A. pittii* and *A. lactucae*), whereas the baumannoferrin *bfnD* is present solely in *A. baumannii*. Similar findings were also reported in Mancilla-Rojano, Flores [63], where *basC* is highly conserved in these three *Acinetobacter* spp. Despite a close phylogenetic relationship, the three genes are absent in *A. nosocomialis*, a pathogen with a mortality rate similar to *A. baumannii* [64], as indicated by the low Blastp coverage and identity. This is likely due to the absence of the clusters. To evaluate this further, we performed the ANI analysis and genomic alignment of the Ab-C98 and the A. nosocomialis strain M2. The analysis demonstrated that *A. nosocomialis* is the closest species to *A. baumannii* (ANI of 91.48%), which is in line with our phylogenetic analysis using the core genes; however, the acinetobactin cluster is completely absent in the *A. nosocomialis* as observed by the genome alignment at the acinetobactin gene locus. This suggests that the acinetobactin cluster is essential in *A. baumannii* virulence as it is highly conserved among the 710 complete genomes analysed. However, it is likely that upon evolutionary transition, *A. nosocomialis* loses the acinetobactin cluster, and the pathogenesis of *A. nosocomialis* may be contributed by alternative virulence mechanisms, such as the type II secretion system [65]. Furthermore, the genome alignment analysis showed that the baumannoferrin cluster is present only in *A. baumannii*, which could likely contribute to the enhanced virulence in this species compared to other non-*A. baumannii Acinetobacter* spp.

## Conclusion

In conclusion, using *G. mellonella* as an *in vivo*-like model, we showed that the community-acquired *A. baumannii* Ab-C98 significantly up-regulated siderophore production pathways and iron transportation during the infection, based on the gene expression analysis from the direct RNA-seq. We confirmed the virulence role of *basC* as observed in previous research studies which used mammalian models. The consistency of our findings with previous studies demonstrated the reliability of *G. mellonella* as an infection model for bacterial virulence studies. Furthermore, this study showed that *A. baumannii* lacked isochorismatase and *bfnD* displayed attenuated virulence and reduced infection, which was previously unknown. *bfnD*, from the baumannoferrin cluster, is specific to *A. baumannii*; however, the high conservation of *basC* (from the acinetobactin cluster) and the isochorismatase in the three closely related pathogenic *Acinetobacter* spp (*A. baumannii*, *A. pittii*, *A. lactucae*) underscores the potential of these two targets as anti-virulence drug targets for reducing the bacterial pathogenesis of the three *Acinetobacter* spp.

## Acknowledgements

The authors thank Professor Bryan W. Davies (The University of Texas at Austin) for providing pAT02 and Nazmul Hasan Muzahid (Monash University Malaysia) for the wild-type *A. baumannii* strains. We thank Monash University Malaysia Genomics Platform for their technical assistance in the MinION sequencing. KET also gratefully acknowledges the receipt of campus scholarships from Monash University Malaysia.

## Funding statement

Not applicable

## Declaration of interest statement

The authors report there are no competing interests to declare.

## Data availability

All Nanopore sequencing data are available at the Sequence Read Archive (SRA, https://www.ncbi.nlm.nih.gov/sra) under the project accession number PRJNA1128375.

## Notes

### Competing Interest Statement

The authors have declared no competing interest.

