## Supplementary for "Transcriptomic Insights into the Virulence of Acinetobacter baumannii During Infection: Role of Iron Uptake and Siderophore Production Genes"

Table S 1: Barcodes for each direct RNA-sequencing library sample.

| Barcode ID | Oligonucleotide name | Oligonucleotide sequences | Sample |
|---|---|---|---|
| BC1 | Oligo A | 5'-/5Phos/GGCTTCTTCTTGCTCTTAGGTAGTAGGTTC-3' | IVB-1 |
| | Oligo B | 5'-GAGGCGAGCGGTCAATTTTCCTAAGAGCAAGAAGAAGCCTTTTTTTTTT-3' | |
| BC2 | Oligo A | 5'-/5Phos/GTGATTCTCGTCTTTCTGCGTAGTAGGTTC-3' | IVV-1 |
| | Oligo B | 5'-GAGGCGAGCGGTCAATTTTCGCAGAAAGACGAGAATCACTTTTTTTTTT-3' | |
| BC3 | Oligo A | 5'-/5Phos/GTACTTTTCTCTTTGCGCGGTAGTAGGTTC-3' | IVB-2 |
| | Oligo B | 5'-GAGGCGAGCGGTCAATTTTCCGCGCAAAGAGAAAAGTACTTTTTTTTTT-3' | |
| BC4 | Oligo A | 5'-/5Phos/GGTCTTCGCTCGGTCTTATTTAGTAGGTTC-3' | IVV-2 |
| | Oligo B | 5'-GAGGCGAGCGGTCAATTTTAATAAGACCGAGCGAAGACCTTTTTTTTTT-3' | |

Table S 2: Oligonucleotides used in the RT-qPCR and gene knockout.

| Primer name | Sequence (5' -3') |
|---|---|

| | |
|---|---|
| RT-qPCR | |
| basE-F | ACACCTGCTGATGAAGTTGC |
| basE-R | GGTTCAGGATTTGGTGCCAT |
| basC-F | AGGTCACGGTTATTGGCTCA |
| basC-R | ATGCAGGGGTAAAGCACTCT |
| iso-F | CTC CGC TCG AAC AAC TCA TG |
| iso-R | TGC CAT ATC GTG CTC TTC TCT |
| bfnD-F | GAC TAA CTT GCA GCA TCC GG |
| bfnD-R | TAG GCC TGT TGA GCA TGT GA |
| bfnA-F | CCA CTA CAT CCT TGG CAA GC |
| bfnA-R | CGC CAC GAT GAC ACT CTT TT |
| deah-F | ACTCACTGGTTGGCTGATGA |
| deah-R | ACTAAGATTTTCGCACGGCC |
| ompW-F | AAGATTTTGGTGTCGCTGGC |
| ompW-R | ACCTAAAGTGTAAACAAACGGGT |
| rpoD-F | AGACCAGAACATCACTTCTCCA |
| rpoD-R | TCACGTGTTACATCAAACTGCT |
| Gene knockout | |
| basC_Up_F | ACAGCAAAATCAACTACTCGAC |
| basC_Up_R | tcggcttttcgccattcgtattgcacgacattgcactccaccgctgatgaACC AAT TCC GGT AAT ATC TAT TTT T |
| basC_Down_F | cgccagaggcgggatgcgaagaatgcgatgccgctcgccagtcgattggcTCT TTC CAG CAC TAT GGT TTA C |
| basC_Down_R | AAT GCA GAA AAT ACC TTG TCC G |
| basC_veri_F | ATA ATG AGT CCG ATC ATT CAT C |
| iso_Up_F | CAA CTC TGG ACA GAT AAT GC |
| iso_ Up _R | tcggcttttcgccattcgtattgcacgacattgcactccaccgctgatgaGCT GTA ACT TGC AAT CTT TGG A |
| iso_Down_F | cgccagaggcgggatgcgaagaatgcgatgccgctcgccagtcgattggcCAA GTA ATC CAA GCA TGG ATA C |
| iso_Down_R | CAT TAT TTT GTA TAA TGG TTA ACA TA |
| iso_veri_F | TTTCACCTGGATCTACTCTC |
| bfnD_Up_F | GCA GCA ACA GCA TCT AGT TA |
| bfnD_Up_R | tcggcttttcgccattcgtattgcacgacattgcactccaccgctgatgaATTTGATAAAACTGGATGTAATTGTC |

| | |
|---|---|
| bfnD_Down_F | cgccagaggcgggatgcgaagaatgcgatgccgctcgccagtcgattggcTTA GCT CCT TTT AAA TTG CAG GT |
| bfnD_Down_R | GGC TTG AGT AAT ACC TTT TTG TA |
| bfnD_veri_F | TGCCATCGACTACGGCAG |
| apmR_F | TCA TCA GCG GTG GAG TGC A |
| apmR_R | GCC AAT CGA CTG GCG AGC |
| apmR_veri_R | GAC GCT ACG GAA GGA GCT |

Table S 3: Quality assessment of the poly(A)-tailed RNA samples using a Nanodrop spectrophotometer.

| Samples | A260/230 | A260/280 | Concentration (ng/µL) |
|---|---|---|---|
| IVB-1 | 2.15 | 2.16 | 104.3 |
| IVV-1 | 2.10 | 2.13 | 178.9 |
| IVB-2 | 2.06 | 2.18 | 152.3 |
| IVV-2 | 2.02 | 2.17 | 118.2 |

Notes: IVB = *In vitro* bacteria from broth culture (no infection control); IVV = *In vivo* bacteria from infected larvae.

Table S 4: General sequencing statistics of the direct RNA-sequencing.

| General summary | Number of reads | |
|---|---|---|
| | raw | filtered |
| Mean read length | 810.1 | 754.6 |
| Mean read quality | 10.8 | 11.3 |
| Median read length | 664.0 | 609.0 |
| Median read quality | 10.9 | 11.4 |
| Number of reads | 2,372,201.0 | 2,224,375.0 |
| Read length N50 | 1,064.0 | 1,010.0 |

| | | |
|---|---|---|
| STDEV read length | 556.4 | 546.0 |
| Total bases | 1,921,648,892.0 | 1,678,564,024.0 |

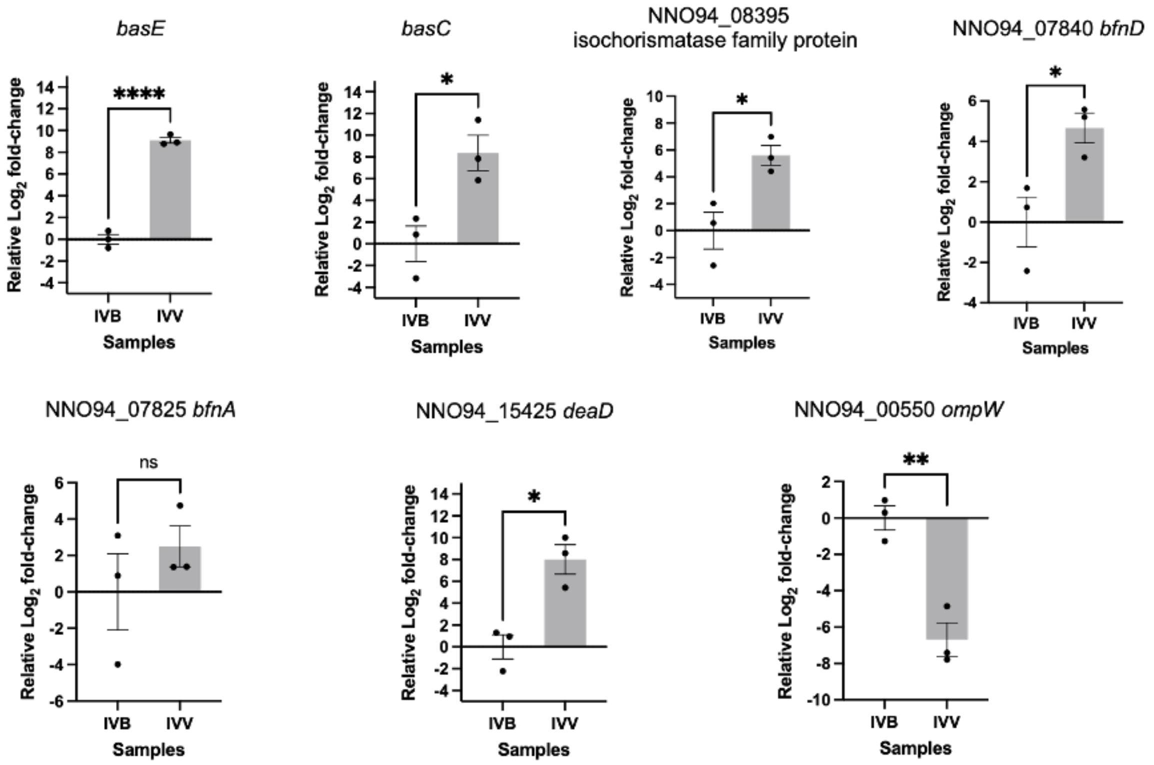


**Figure S 1: RT-qPCR of the 7 shortlisted differentially expressed genes.** The fold-change values represent the $log_2$-scaled relative to the no infection control. Three independent biological replicates with three technical replicates each were performed. The error bars represent the mean± SEM. Asterisks on the qPCR values indicate a significant difference between the infection sample and non-infection control Ab-C98, using an unpaired two-tailed Student’s t-test. p<0.05 (*), p<0.01 (**), p<0.001 (***), p<0.0001 (****), ns (non-significant). IVB = no infection control (Ab-C98 broth culture), IVV = infection sample (Ab-C98 isolated from infected *G. mellonella*)

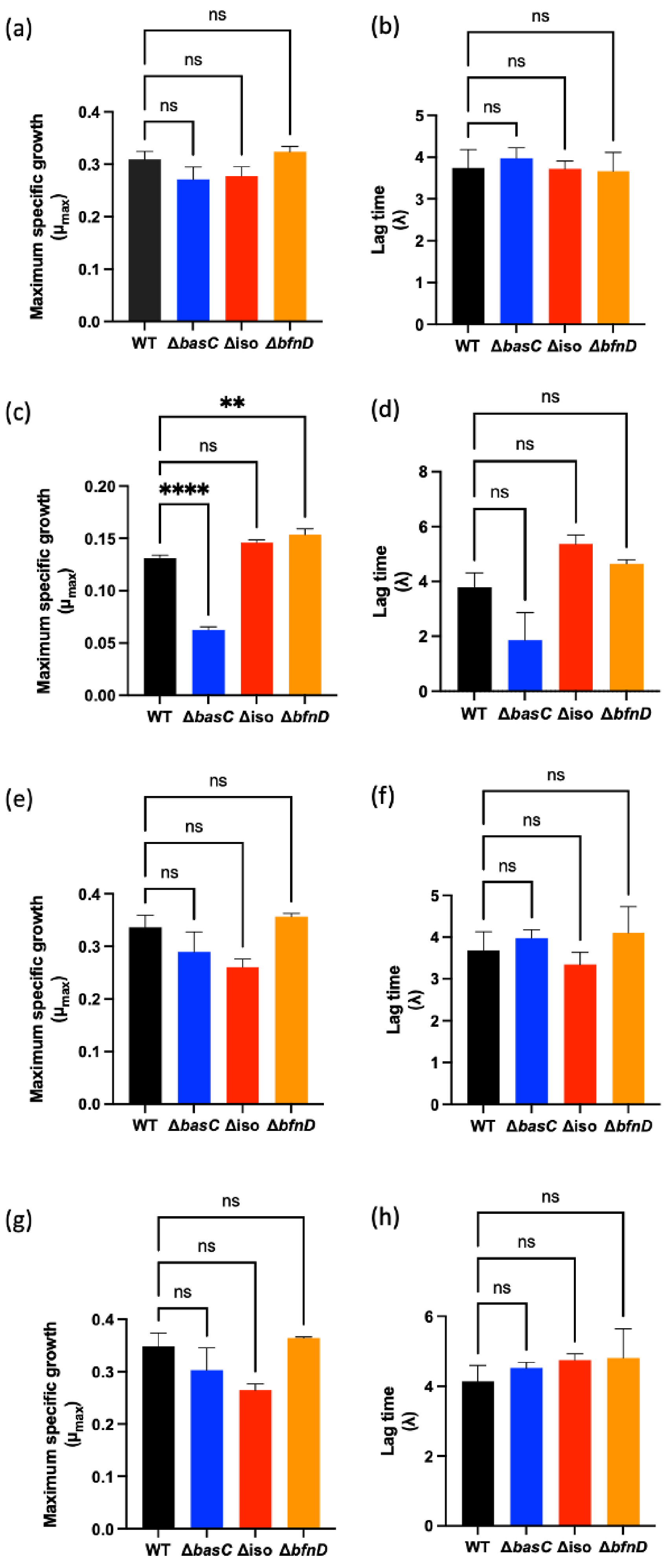
(a)
(b)
(c)
(d)
(e)
(f)
(g)
(h)
ns
**
****
Maximum specific growth
(μmax)
Lag time
(λ)
WT
ΔbasC
Δiso
ΔbfnD
0.0
0.1
0.2
0.3
0.4
0.00
0.05
0.10
0.15
0.20
0
1
2
3
4
5
6
8

**Figure S 2: Growth kinetics of *A. baumannii* strain C98 and its isogenic mutant derivatives** under the conditions of (a-b) MHB culture medium only, (c-d) 200 µM BIP (iron-depleted), (e-f) 200 µM FeCl3 (iron-rich), and (g-h) 200 µM BIP and 200 µM FeCl3. Three independent biological replicates with three technical replicates each were performed. Asterisks indicate a statistical significance between the knockout mutants and the wild-type. $p<0.05$ (*), $p<0.01$ (**), $p<0.001$ (***), $p<0.0001$ (****), ns (non-significant).

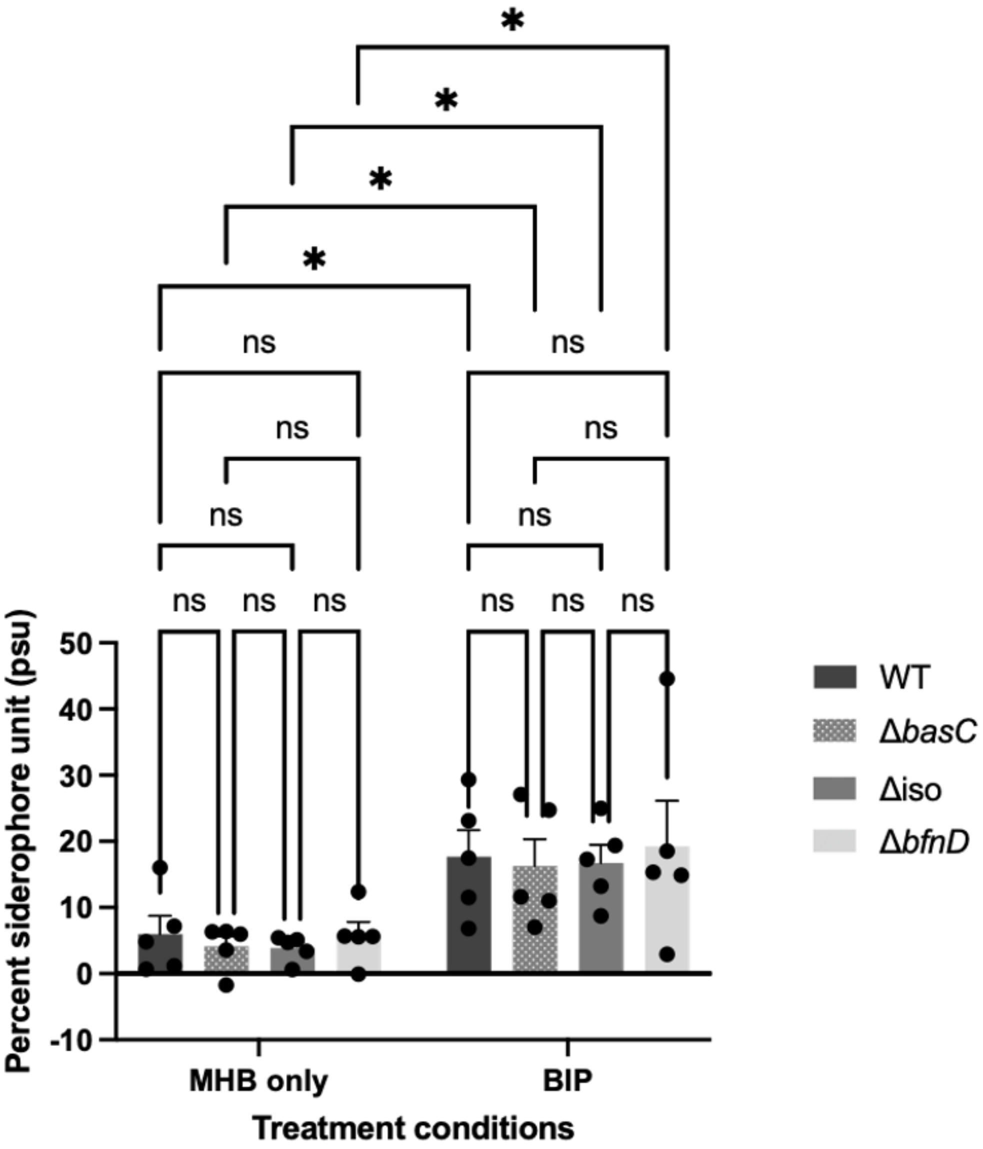


| | MHB | | | | BIP | | | |
|---|---|---|---|---|---|---|---|---|
| | WT | Δ*basC* | Δ*iso* | Δ*bfnD* | WT | Δ*basC* | Δ*iso* | Δ*bfnD* |
| Average psu | 5.965 | 4.104 | 3.858 | 5.830 | 17.657 | 16.295 | 16.707 | 19.243 |

**Figure S 3: Total siderophore activity of wild-type Ab-C98 and its mutant derivatives using Chrome Azurol S (CAS) microplate method.** The overall production of siderophores was expressed in percent siderophore unit (psu) normalised to the reference control (uninoculated broth medium). The average psu of each strain was summarised in the table. Five independent biological replicates with three technical replicates each were performed. The error bars represent the mean± SEM. The statistical significance difference between the strains and the two treatment conditions was analysed via two-way ANOVA with Tukey's test. A *p*-value $< 0.05$ (*) indicates significance.
